# Dietary Protein on Growth Performance, Serum Biochemical Index and offspring of Hu ewes in late pregnancy

**DOI:** 10.1101/2020.08.04.235853

**Authors:** Xin Wang, Qiye Wang, Yancan Wang, Chunpeng Dai, Jianzhong Li, Pengfei Huang, Yali Li, Xueqin Ding, Jing Huang, Tarique Hussain, Huansheng Yang

## Abstract

The purpose of this study was to explore the effects of ewe growth performance, serum biochemical indicators and lamb growth and development at different protein levels in late pregnancy. A total of 15 pregnant ewes (46.4 ± 1.38kg initial BW) were assigned to 3 groups with 5 ewes in each group in a randomized block design. P1, P2 and P3 of three groups were fed diets with different levels of crude protein at 11.25%, 12.5% and 13.75% (low, medium and high) respectively, while the dietary energy levels remained unchanged. Choose ewes with the same body condition from 90 days of pregnancy to different protein diets during delivery, and feed the same diets during lactation. New-born lambs were left to suckle their dams freely for the first three days and were kept with their dams till weaning at 60 days of age. Dietary protein levels during late gestation showed no significant effect on the weight gain of ewes and their offspring in the late pregnancy (*P* > 0.05) but had significant effects on blood urea nitrogen (BUN), glucose (GLU), C-reactive protein (CRPL3) and ammonia nitrogen (NH3L) before parturition, and on triglyceride (TG) in lactating ewes. Dietary protein levels during late gestation had significant effects on birth height, body length, chest depth, chest circumference, straight crown hip length and curved crown hip length of lambs (*P* < 0.05) as well as on weaning body length, chest circumference, head width and head length (*P* < 0.05). The results showed that although different dietary protein levels during late gestation have no significant effects on growth performance of ewes, singleton, twins and triplets, it affects lambs’ body shape and ewes’ metabolism. Therefore, the optimum protein concentration for the growth of ewes and subsequent lambs in this experiment is 11.25%.

## 1 Introduction

The nutritional status of ewes during pregnancy is crucial because it has a great impact on the development of the foetus, survival and performance of subsequent lambs by affecting the fecundity and the colostrum production of ewes (Bell, 1995). In the late trimester, the mother’s metabolism is mainly focused on providing energy for the development and growth of the foetus (Mohammadi et al., 2016), and the animal regulates carbohydrate, protein and lipid metabolism (Bell et al., 1995) to maintain its homeostasis and deliver nutrients to the pregnant uterus and lactating breast (Joy et al., 2014). During pregnancy, if malnourished, the female animal mobilises a large amount of muscle and fat to maintain normal foetal development. Chronic malnutrition can lead to wasting syndrome. The energy and protein deficiency of ewes will not only harm the ewe but also affect the survival and production performance of lambs (González-Recio et al., 2012). Maternal malnutrition and over-nutrition can significantly affect foetal growth and development (Caton et al., 2010), and the development and production performance of lambs through prenatal programs. Providing reasonable nutrition according to the maternal and foetal requirements is important to improve the reproductive performance of females.

Protein, as the nutrient of life, plays an important role in the growth and reproduction of organisms. Ocak et al. (2005) investigated the effect of feeding high dietary protein levels during late gestation to ewes on colostrum yield and lamb survival rate and concluded that an excess of CP in the diet caused a decrease in colostrum yield and lamb survival. This was due to the increased lamb birth weight resulting in a higher number of lamb dystocia. Compared to lambs born as singleton, twins and triplets have a lower birth weight and lower weaning weight (Gardner et al., 2007). Also, lambs born as triplets but reared as twins have been found to have lower weaning weight than lambs born and reared as twins (Morris and Kenyon, 2004). An increase in litter size per ewe should offer greater financial gains, but is also associated with high mortality rates, and is consequently a potential welfare concern. For this reason, there is a growing interest in optimal feeding regimens for these prolific ewes given their higher theoretical nutritional requirements in the last trimester of gestation. Consequently, the CP level of diets fed to pregnant ewes during this critical period of gestation and ensuing lactation has been a concern for livestock scientists. It becomes important to determine the pre- and post-partum dietary protein levels that can maximise the production potential of ewes and the performance of their lambs. Therefore, the purpose of this study was to investigate the effects of different protein levels on the growth and development of ewes and lambs of a local Chinese sheep breed during late pregnancy and whether this effect extends to lactation.

## 2 Materials and methods

### 2.1 Ethics statement

All experimental procedures related to the animals were performed according to protocols approved by the Animal Care Advisory Committee of Hunan Normal University, Changsha, Hunan, China.

### 2.2 Diets, animals, and e, xperimental procedures

This experiment was performed at the Hubei Zhiqinghe griculture and Animal Husbandry, the area is mainly subtropical monsoon climate, and Hu sheep has a strong adaptability to this climate. A total of 15 pregnant ewes (46.4 ± 1.38kg initial BW) were assigned to 3 groups with 5 ewes in each group in a randomized block design. Determine the gestation time of each ewes and whether they are pregnant by performing simultaneous estrus and B-ultrasound monitoring on the ewes. P1, P2 and P3 of three groups were fed diets with different levels of crude protein at 11.25%, 12.5% and 13.75% (low, medium and high) respectively, while the dietary energy levels remained unchanged. The experimental diet of ewes in the late pregnancy is shown in Table 1 and the experimental diet of lambs is shown in Table 2. Diets were provided daily at 07:00 and 17:00 h. Feed was formulated as a mixed diet. Each treatment was fed in individual pens (length × width × height = 5.0 × 2.5 × 1.0 m) with individual feeding and automatic water. Ewe and lamb feed was designed according to NRC2007, ewe feed was designed according to the diet formula of 50kg ewe weight and 1.45kg daily gain, lamb feed was designed according to the diet formula of 20kg lamb weight and 200g daily gain, protein and other nutrients met the requirements of all test ewes. The pregnancy period and the whole lactation period of ewes were weighed every ten days on a digital multi-function platform scale. Record the lamb number of ewes, weigh the lambs within 24 hours after birth, and weigh the lambs every ten days until they are weaned. Each group of ewes and their lambs were housed in a partly enclosed yard. New-born lambs were left to suckle their dams freely for the first three days and were kept with their dams till weaning at 60 days of age.

**Table 1.** Dietary composition of ewes with different nutritional levels in the third trimester of pregnancy (air-dried basic%)

| Items | DM | Groups |  |  |
| --- | --- | --- | --- | --- |
|  |  | P1 | P2 | P3 |
| Ingredients, % |  |  |  |  |
| Corn silage | 34.00 | 30 | 30 | 30 |
| Peanut seedling | 91.30 | 30 | 30 | 30 |
| Corn | 86.00 | 27.6 | 25 | 22.4 |
| Wheat bran | 87.00 | 6 | 5.4 | 4.8 |
| Soybean meal | 89.00 | 3.4 | 6.6 | 9.8 |
| NaCl | 100.00 | 0.3 | 0.3 | 0.3 |
| NaHCO3 | 100.00 | 0.5 | 0.5 | 0.5 |
| Premix<sup>1</sup> | 100.00 | 2.2 | 2.2 | 2.2 |
| Total |  | 100 | 100 | 100 |
| Nutrient levels<sup>2</sup>, DM |  |  |  |  |
| ME, MJ/kg |  | 10 | 10 | 10 |
| CP, % |  | 11.25 | 12.5 | 13.75 |

Notes. <sup>1</sup>Premix provides the following per kg: Vitamin A, 120KIU; Vitamin D<sub>3</sub>, 60KIU; Vitamin E, 200mg; Cu, 0.15g; Fe, 1g; Zn, 1g; Mn, 0.5g; I, 15mg; Se, 5mg; Co, 2.5mg; Ca, 20g; NaCl, 100-250g; P, 10g. <sup>2</sup>Except for metabolic energy was calculated value, the rest were measured values

**Table 2.** Dietary composition of lambs with different nutrition levels during lactation (air-dried basic%)

| items | DM | group |  |  |
| --- | --- | --- | --- | --- |
|  |  | P1 | P2 | P3 |
| Ingredients |  |  |  |  |
| Corn silage | 23.00 | 20.00 | 20.00 | 20.00 |
| Peanut seedling | 91.30 | 30.00 | 30.00 | 30.00 |
| Corn | 86.00 | 24.90 | 22.93 | 20.96 |
| Wheat bran | 87.00 | 3.95 | 3.52 | 3.09 |
| Soybean meal | 89.00 | 18.15 | 20.55 | 22.96 |
| Premix | 100.00 | 3.00 | 3.00 | 3.00 |

Notes. <sup>1</sup>Premix provides the following per kg: Vitamin A, 120KIU; Vitamin D<sub>3</sub>, 60KIU; Vitamin E, 200mg; Cu, 0.15g; Fe, 1g; Zn, 1g; Mn, 0.5g; I, 15mg; Se, 5mg; Co, 2.5mg; Ca, 20g; NaCl, 100-250g; P, 10g. <sup>2</sup>Except for metabolic energy was calculated value, the rest were measured values

### 2.3 Growth performance

During the experiment, all the lambs and ewes were weighed individually in the morning on an empty stomach. The ewes were weighed before the beginning of the experiment and a week before parturition, and the lambs were weighed on the 0th day after birth, then weighed every 10 days. The average daily gain (ADG) was calculated on the basis of the average of 3 measurements for individual ADG (Yin et al., 2001). The lamb’s litter weight is the average weight of all lambs born to each ewe.

### 2.4 Blood biochemical indexes

Venous blood samples were taken from the neck of five ewes in each group at 140 days of pregnancy, 145 days of pregnancy, 1 day and 20 days after delivery before morning feeding. Venous blood was taken from the neck of the lamb on the 60th day of lactation and samples were collected with 5ml vacuum tube (Hunan Changsha Yiqun Medical equipment Co., Ltd.). The supernatant was stored at-80 °C for biochemical parameters analysis after centrifugation (3500 RMP par 4 °C, 15min). A TBA-120FR Automatic Biochemistry Radiometer (Hitachi) was used to measure the concentrations of serum biochemical indices (Chen et al., 2019; Yin et al., 2018).

### 2.5 Milk composition

The milk samples were hand milked both in the morning and afternoon (8:00, 17:00), and be collected continuously on the first and second day of lactation, 20 ml each time, mixed and preserved at –20°C with the antimicrobial preservative Bronopol (2-bromo-2-nitro-1,3-propanediol). They were analysed for fat, protein, lactose content by NIRS (MilkoScan FT6000) and SCC by a fluorimetric method (Fossomatics) (both devices from Foss Electric A/S, Hillered, Denmark) (Madeline Koczura et al., 2019).

### 2.6 Body measurements of lambs

The body size of lambs was measured at 0 and 60 days after birth. The indexes and methods were as follows (Kabir et al., 2014):

- Body height: The vertical distance from the shoulder to the ground
- Body length: The distance from the shoulder to the end of the ischial tuberosity
- Chest depth: The vertical distance from the nail to the lower edge of the sternum
- Chest circumference: The length of the posterior edge of the scapula around the chest
- Abdominal circumference: The vertical circumference of the largest part of the abdomen
- Head width: Length of transverse muscle lines on both sides of the head
- Head length: The vertical distance from the top of the head to the chin
- Straight crown hip length: The vertical distance from the forehead to the end of the coccyx
- Curved crown hip length: The distance between the end of the forehead along the back and the end of the tailbone

### 2.7 Statistical analysis

Excel (Microsoft) was used for preliminary processing of test data processing. Spss 18.0 software (SPSS) was used for data variance analysis of data, and Duncan’s tests were used for multiple comparisons among different groups. The final test results were presented with mean ± SEM. A p-value of <0.05 was significant.

## 3 Results and analysis

### 3.1 Growth performance of ewes and lambs

Table 3 shows that the initial weight, final weight, total gain and ADG did not have a significant difference in the different protein levels of ewes (*P* > 0.05), but the ADG in the low protein group was 28.14% and 32.79% higher than in the medium protein group and high protein group. In Table 4, there was no significant difference in the body weight of singletons, twins and triplets born by ewes at different protein levels (*P* > 0.05). The litter birth weight, litter weight per 10 days and litter weight gain per 10 days in each group have been presented in Table 5. There was no significant difference in litter birth weight, 10-day litter weight gain or 10-day litter weight among the three groups (*P* > 0.05). In the birth weight of lambs there were no significant differences in body weight and body weight gain per 10 days during lactation (*P* > 0.05; Table 6).

**Table 3.** effects of different protein levels on growth performance of ewes in late gestation period.

| Items | Groups |  |  | SEM | P-value |
| --- | --- | --- | --- | --- | --- |
|  | P1 | P2 | P3 |  |  |
| Initial weight, kg | 48.55 | 44.71 | 48.9 | 3.22 | 0.382 |
| Final weight, kg | 58.06 | 51.74 | 57.13 | 3.42 | 0.179 |
| Total Gain, kg | 9.51 | 7.03 | 8.23 | 1.34 | 0.223 |
| ADG, g | 221.16 | 163.49 | 191.40 | 31.22 | 0.223 |

Notes: Low protein (P1, 11.25%), medium protein (P2, 12.5), high protein (P3,13.75). ADG – average daily gain. SEM, standard error of the mean. N = 5.

**Table 4.** The weight of singleton, twins and triplet lambs.

| Items (kg) | Group |  |  | SEM | P-value |
| --- | --- | --- | --- | --- | --- |
|  | P1 | P2 | P3 |  |  |
| Singleton |  |  |  |  |  |
| Initial weight | 5.1 | 3.9 | 4 | _____ | _____ |
| Final weight | 23.7 | 18.7 | 20.7 | _____ | _____ |
| Total weight gain | 18.6 | 14.8 | 16.7 | _____ | _____ |
| Twins |  |  |  |  |  |
| Initial weight | 3.2 | 3.4 | 3.6 | 0.955 | 0.407 |
| Final weight | 13.9 | 14.05 | 16.38 | 1.816 | 0.197 |
| Total weight gain | 10.7 | 10.65 | 12.78 | 1.82 | 0.196 |
| Triplets |  |  |  |  |  |
| Initial weight | 3.1 | 3.4 | 2.7 | 1.883 | 0.232 |
| Final weight | 13 | 13.67 | 13.63 | 0.082 | 0.923 |
| Total weight gain | 9.9 | 10.27 | 10.93 | 0.218 | 0.81 |

Notes: Low protein (P1, 11.25%), medium protein (P2, 12.5%), high protein (P3,13.75%). SEM, standard error of the mean.

**Table 5.** Effects of different protein levels in late gestation period on litter weight gain of lambs.

| Items | Groups |  |  | SEM | P value |
| --- | --- | --- | --- | --- | --- |
|  | P1 | P2 | P3 |  |  |
| Litter weight, kg |  |  |  |  |  |
| d 0 | 6.76 | 6.86 | 6.72 | 1.17 | 0.992 |
| d 10 | 9.6 | 9.7 | 10.32 | 1.79 | 0.661 |
| d 20 | 11.96 | 12.42 | 13.3 | 1.52 | 0.677 |
| d 30 | 15.28 | 15.8 | 17.12 | 1.98 | 0.642 |
| d 40 | 19.28 | 19.66 | 21.78 | 2.5 | 0.575 |
| d 50 | 24.04 | 23.92 | 26.62 | 3.57 | 0.701 |
| d 60 | 29.26 | 28.76 | 31.98 | 4.68 | 0.765 |
| Litter weight gain, kg |  |  |  |  |  |
| d 0-10 | 2.84 | 2.84 | 3.6 | 0.5 | 0.259 |
| d 10-20 | 2.36 | 2.72 | 2.98 | 0.36 | 0.268 |
| d 20-30 | 3.32 | 3.38 | 3.82 | 0.63 | 0.681 |
| d 30-40 | 4 | 3.86 | 4.66 | 0.64 | 0.440 |
| d 40-50 | 4.76 | 4.26 | 4.84 | 1.19 | 0.872 |
| d 50-60 | 5.22 | 4.84 | 5.36 | 1.36 | 0.925 |
| Total gain | 22.5 | 21.9 | 25.26 | 3.58 | 0.618 |

Notes: Low protein (P1, 11.25%), medium protein (P2, 12.5%), high protein (P3,13.75%).
SEM, standard error of the mean. N = 5.

**Table 6.** Effects of different protein levels on the growth performance of lambs.

| Items | Groups |  |  | SEM | P-value |
| --- | --- | --- | --- | --- | --- |
|  | P1 | P2 | P3 |  |  |
| Body weight, kg |  |  |  |  |  |
| d 0 | 3.38 | 3.43 | 3.36 | 0.27 | 0.966 |
| d 10 | 4.8 | 4.85 | 5.16 | 0.48 | 0.718 |
| d 20 | 5.98 | 6.21 | 6.65 | 0.7 | 0.627 |
| d 30 | 7.64 | 7.9 | 8.56 | 0.95 | 0.612 |
| d 40 | 9.64 | 9.83 | 10.89 | 1.17 | 0.523 |
| d 50 | 12.02 | 11.96 | 13.31 | 1.26 | 0.489 |
| d 60 | 14.63 | 14.38 | 15.99 | 1.44 | 0.492 |
| Body weight gain, kg |  |  |  |  |  |
| d 0-10 | 1.42 | 1.42 | 1.8 | 0.25 | 0.242 |
| d 10-20 | 1.18 | 1.36 | 1.49 | 0.27 | 0.532 |
| d 20-30 | 1.66 | 1.69 | 1.91 | 0.31 | 0.682 |
| d 30-40 | 2 | 1.93 | 2.33 | 0.29 | 0.362 |
| d 40-50 | 2.38 | 2.13 | 2.42 | 0.25 | 0.471 |
| d 50-60 | 2.61 | 2.42 | 2.68 | 0.39 | 0.793 |
| Total gain | 11.25 | 10.95 | 12.63 | 1.22 | 0.354 |

Notes: Low protein (P1, 11.25%), medium protein (P2, 12.5%), high protein (P3,13.75%).
SEM, standard error of the mean. N = 5.

### 3.2 Blood biochemical indexes of ewes and lambs

Dietary treatments with different protein levels showed significant differences in BUN, GLU, CRPL3 and NH3L in ewes before delivery (*P* < 0.05; Table 7). Serum TG content in the P2 group was significantly greater (*P* < 0.05, Table 8) than P1 and P3 groups. No significant difference in biochemical indexes of weaned lambs was observed among the three groups (*P*>0.05; Table 9).

**Table 7.** Blood biochemical indexes of ewes in late pregnancy before delivery.

| Items | Groups |  |  | SEM | *P* value |
| --- | --- | --- | --- | --- | --- |
|  | P1 | P2 | P3 |  |  |
| TP, g/L | 59.53 | 64.14 | 63 | 1.095 | 0.218 |
| ALB, g/L | 31.77 | 31.27 | 32.21 | 0.504 | 0.756 |
| ALT, U/L | 14.07 | 13.83 | 12.97 | 0.637 | 0.773 |
| AST, U/L | 69.78 | 74.9 | 68.82 | 2.018 | 0.425 |
| AST/ALT | 4.96 | 5.4 | 4.79 | 0.167 | 0.313 |
| LDH, U/L | 341.7 | 304.6 | 314.36 | 12.883 | 0.501 |
| BUN, mmol/L | 3.44<sup>b</sup> | 5.17<sup>a</sup> | 5.36<sup>a</sup> | 0.256 | 0.001 |
| CREA, μmol/L | 54.4 | 66.33 | 59.45 | 2.052 | 0.063 |
| GLU, mmol/L | 1.11<sup>b</sup> | 1.42<sup>a</sup> | 1.51<sup>a</sup> | 0.067 | 0.029 |
| TG, mmol/L | 0.43 | 0.43 | 0.51 | 0.02 | 0.181 |
| CHOL, mmol/L | 1.86 | 1.93 | 1.81 | 0.041 | 0.484 |
| HDL, mmol/L | 1.09 | 1.06 | 1.06 | 0.025 | 0.926 |
| LDL-C3, mmol/L | 0.6 | 0.68 | 0.6 | 0.024 | 0.312 |
| CRPL3, mmol/L | 4.32<sup>b</sup> | 4.31<sup>b</sup> | 4.34<sup>a</sup> | 0.004 | 0.006 |
| LACT, mmol/L | 3.92 | 4.76 | 4.42 | 0.152 | 0.082 |
| NH3L, μmol/L | 235.37<sup>b</sup> | 245.47<sup>ab</sup> | 265.87<sup>a</sup> | 4.619 | 0.017 |

Notes: Low protein (P1, 11.25%), medium protein (P2, 12.5%), high protein (P3,13.75%).
TP, total protein; ALB, albumin ; ALT, alanine aminotransferase; AST, aspartate aminotransferase; LDH, lactic dehydrogenase; BUN, blood urea nitrogen; CREA, creatinine; GLU, glucose; TG, triglyceride; CHOL, cholesterol; HDL, high density lipoprotein; LDL-C3, low density lipoprotein; CRPL3, C-Reactive protein; LACT, lecithin-cholesterol acyltransferase; NH3L, ammonia.
SEM, standard error of the mean. N = 5.

**Table 8.** Blood biochemical indexes of ewes in suckling period before weaning.

| Items | Groups |  |  | SEM | *P* value |
| --- | --- | --- | --- | --- | --- |
|  | P1 | P2 | P3 |  |  |
| TP, g/L | 68.07 | 68.49 | 72.46 | 1.153 | 0.25 |
| ALB, g/L | 31.61 | 32.81 | 33.42 | 0.623 | 0.518 |
| ALT, U/L | 22.66 | 23 | 22.94 | 1.258 | 0.994 |
| AST, U/L | 95.4 | 109.95 | 102.23 | 4.056 | 0.397 |
| AST/ALT | 4.26 | 4.62 | 4.48 | 0.245 | 0.855 |
| LDH, U/L | 523.86 | 505.75 | 509.4 | 18.776 | 0.931 |
| BUN, mmol/L | 3.79 | 4.03 | 3.94 | 0.278 | 0.949 |
| CREA, μmol/L | 46.43 | 52.63 | 49.11 | 1.271 | 0.414 |
| GLU, mmol/L | 3.09 | 3.27 | 3.22 | 0.088 | 0.748 |
| TG, mmol/L | 0.22<sup>b</sup> | 0.29<sup>a</sup> | 0.22<sup>b</sup> | 0.013 | 0.036 |
| CHOL, mmol/L | 1.99 | 2.22 | 2.39 | 0.082 | 0.071 |
| HDL, mmol/L | 1.55 | 1.68 | 1.81 | 0.07 | 0.171 |
| LDL-C3, mmol/L | 0.4 | 0.5 | 0.52 | 0.027 | 0.193 |
| CRPL3, mmol/L | 4.35 | 4.36 | 4.37 | 0.005 | 0.455 |
| LACT, mmol/L | 4.39 | 5.17 | 4.82 | 0.258 | 0.533 |
| NH3L, μmol/L | 195.49 | 201.89 | 185.29 | 5.471 | 0.455 |

Notes: Low protein (P1, 11.25%), medium protein (P2, 12.5%), high protein (P3,13.75%).
TP, total protein; ALB, albumin ; ALT, alanine aminotransferase; AST, aspartate aminotransferase; LDH, lactic dehydrogenase; BUN, blood urea nitrogen; CREA, creatinine; GLU, glucose; TG, triglyceride; CHOL, cholesterol; HDL, high density lipoprotein; LDL-C3, low density lipoprotein; CRPL3, C-Reactive protein; LACT, lecithin-cholesterol acyltransferase; NH3L, ammonia.
SEM, standard error of the mean. N = 5.

**Table 9.** Effect of maternal protein level on blood biochemical indexes of weaned lambs.

| Items | Groups |  |  | SEM | P value |
| --- | --- | --- | --- | --- | --- |
|  | P1 | P2 | P3 |  |  |
| TP, g/L | 54.32 | 49.47 | 54.62 | 1.075 | 0.128 |
| ALB, g/L | 32.57 | 30.58 | 33.17 | 0.756 | 0.366 |
| ALT, U/L | 13.52 | 12.72 | 14.73 | 0.687 | 0.511 |
| AST, U/L | 78.33 | 71.67 | 77.4 | 3.383 | 0.702 |
| AST/ALT | 5.99 | 5.76 | 7.71 | 0.736 | 0.526 |
| LDH, U/L | 486.67 | 471.67 | 568 | 26.45 | 0.294 |
| BUN, mmol/L | 2.72 | 2.72 | 2.95 | 0.237 | 0.909 |
| CREA, μmol/L | 39 | 37.17 | 37.67 | 1.796 | 0.921 |
| GLU, mmol/L | 3.6 | 3.18 | 3.48 | 0.098 | 0.205 |
| TG, mmol/L | 0.36 | 0.36 | 0.28 | 0.02 | 0.154 |
| CHOL, mmol/L | 1.37 | 1.26 | 1.41 | 0.065 | 0.67 |
| HDL, mmol/L | 0.88 | 0.85 | 0.96 | 0.048 | 0.653 |
| LDL-C3, mmol/L | 0.37 | 0.32 | 0.39 | 0.028 | 0.674 |
| CRPL3, mmol/L | 4.35 | 4.34 | 4.34 | 0.006 | 0.837 |
| LACT, mmol/L | 4.23 | 4.68 | 4.98 | 0.284 | 0.62 |
| NH3L, μmol/L | 193.67 | 193.23 | 214.93 | 5.839 | 0.232 |

Notes: Low protein (P1, 11.25%), medium protein (P2, 12.5%), high protein (P3,13.75%).
TP, total protein; ALB, albumin ; ALT, alanine aminotransferase; AST, aspartate aminotransferase; LDH, lactic dehydrogenase; BUN, blood urea nitrogen; CREA, creatinine; GLU, glucose; TG, triglyceride; CHOL, cholesterol; HDL, high density lipoprotein; LDL-C3, low density lipoprotein; CRPL3, C-Reactive protein; LACT, lecithin-cholesterol acyltransferase; NH3L, ammonia.
SEM, standard error of the mean. N = 5.

### 3.3 Body size of the lamb

There were significant differences in body height, chest depth, chest circumference, straight crown hip length and curved crown hip length among lambs in the born body measurements (*P* < 0.05). The body height, chest depth and chest circumference of the P1 group were significantly higher than those of P2 and P3 group, and the straight crown hip length and curved crown hip length of the P3 group were significantly higher than those of the P1 and P2 groups (*P* < 0.05; Table 10).

**Table 10.** Effect of dietary protein of ewes during late gestation on body size of their offspring.

| Items | Groups |  |  | SEM | *P*-value |
| --- | --- | --- | --- | --- | --- |
|  | P1 | P2 | P3 |  |  |
| Born body measurements, cm (Within 24 h) |  |  |  |  |  |
| Body height | 30.00<sup>a</sup> | 28.80<sup>b</sup> | 29.31<sup>ab</sup> | 0.18 | 0.037 |
| Body length | 31.92 | 30.65 | 31.75 | 0.24 | 0.055 |
| Chest deep | 14.25<sup>a</sup> | 13.50<sup>b</sup> | 13.86<sup>ab</sup> | 0.11 | 0.021 |
| Chest circumference | 35.17<sup>a</sup> | 33.28<sup>b</sup> | 35.17<sup>a</sup> | 0.34 | 0.022 |
| Abdominal girth | 32.50 | 30.45 | 32.44 | 0.46 | 0.093 |
| Head width | 6.42 | 6.20 | 6.42 | 0.05 | 0.071 |
| Head length | 10.75 | 10.53 | 10.67 | 0.06 | 0.324 |
| Crown length of straight | 43.75<sup>ab</sup> | 42.60<sup>b</sup> | 45.03<sup>a</sup> | 0.36 | 0.009 |
| Curved crown hip length | 50.92<sup>a</sup> | 48.28<sup>b</sup> | 51.78<sup>a</sup> | 0.49 | 0.003 |
| Weaning body measurements, cm (60d) |  |  |  |  |  |
| Body height | 35.92 | 35.28 | 36.28 | 0.28 | 0.311 |
| Body length | 51.83<sup>ab</sup> | 47.06<sup>b</sup> | 53.00<sup>a</sup> | 1.10 | 0.047 |
| Chest deep | 25.33 | 24.14 | 25.06 | 0.27 | 0.166 |
| Chest circumference | 56.96<sup>ab</sup> | 53.86<sup>b</sup> | 57.47<sup>a</sup> | 0.66 | 0.037 |
| Abdominal girth | 59.17 | 55.75 | 58.75 | 0.67 | 0.069 |
| Head width | 8.21<sup>ab</sup> | 8.02<sup>b</sup> | 8.52<sup>a</sup> | 0.07 | 0.005 |
| Head length | 16.00<sup>ab</sup> | 15.57<sup>b</sup> | 16.39<sup>a</sup> | 0.12 | 0.007 |
| Crown length of straight | 67.67 | 65.11 | 68.61 | 0.80 | 0.154 |
| Curved crown hip length | 82.58 | 80.06 | 85.72 | 1.03 | 0.053 |

Notes: Low protein (P1, 11.25%), medium protein (P2, 12.5%), high protein (P3,13.75%).
SEM, standard error of the mean. N= 5.

For the weaning body size of lambs, significant differences were observed in body length, chest circumference, head width and head length (*P* < 0.05). The size of the P3 group was significantly higher than that of the P1 and P2 groups (*P* < 0.05, Table 10).

### 3.4 Milk components in ewes

The results of protein composition analysis of ewe milk in late pregnancy have been shown in Table 11. Different protein levels had no significant effect on the milk fat rate, milk protein rate, lactose rate, fat-free dry matter, total dry matter and urea nitrogen (*P* > 0.05).

**Table 11.** Analysis results of milk composition of ewe protein level in late gestation.

| Items | Groups |  |  | SEM | *P*-value |
| --- | --- | --- | --- | --- | --- |
|  | P1 | P2 | P3 |  |  |
| Butter fat rate, % | 5.79 | 6.2 | 8.22 | 0.725 | 0.325 |
| Milk protein, % | 7.53 | 7.72 | 8.19 | 0.208 | 0.394 |
| Rate of lactose, % | 5.26 | 5.11 | 5.09 | 0.093 | 0.795 |
| Remove fat and dry substances, % | 13.83 | 14.35 | 14.91 | 0.315 | 0.374 |
| The total dry matter, % | 19.6 | 21.27 | 23.01 | 0.679 | 0.112 |
| Urea nitrogen, mg/dL | 22.5 | 26.46 | 26.22 | 1.579 | 0.558 |

Notes: Low protein (P1, 11.25%), medium protein (P2, 12.5%), high protein (P3,13.75%). SEM, standard error of the mean. N = 5.

## 4 Discussion

The change in body weight of ewes during pregnancy is an important index to assess the effect of nutrition intake during pregnancy. When the nutrition received is insufficient, the nutrients obtained by ewes will be used in the growth and development of embryos, thus reducing the body weight gain of ewes. In this experiment, diets with different protein levels had no effect on the growth performance of ewes, and the postpartum weight of ewes was higher than that at 90 days of pregnancy, which suggests that all the ewes were in good condition, and the diets met their requirements for gestation and lactation. This implies that the diets supported foetal development and lactation without the ewes metabolising their body reserves. So, the low protein content of 11.25% provided in this experiment can meet the protein needs of normal growth in ewes during late pregnancy.

In this experiment, no significant difference was observed in the effect of different protein levels on the growth performance of lambs, which was consistent with the results of Ocak, N (2005), who concluded that there was no difference in body weight of lambs weaned with different protein levels. Aziz and Al-Dabbagh (2008) pointed out that diets with high nutritional levels had no effect on the BW of lambs and weaned lambs at the end of pregnancy. Voso (2014) showed that high levels of protein can increase the daily gain of lambs. The overall weight gains with low protein, mid-level protein and high protein intake were 11.52 kg, 10.36 kg and 12.31 kg respectively in this experiment. This is consistent with Voso’s conclusion. Although the final body weight of the lambs in the high protein group was higher than that in the other two groups, there was no difference in weight among the three groups. In the lamb samples selected in this experiment, there was no difference in body weight between twins and triplets. McGovern et al also emphasized this point (2015). Changing the protein level of twin ewes in the late pregnancy will not affect the birth weight or organ weight of newborn lambs.

Sheep body type index reflects the growth and development of sheep and indirectly reflects the development of sheep tissues and organs. Although body weight is one of the indicators reflecting the growth and development of animals, it cannot reflect the development of specific tissues and organs of animals. Therefore, it is important to determine the body size of lambs to evaluate the growth and development of lambs. Adequate and reasonable maternal nutrition is necessary for the healthy development of the foetus. The foetal period is also a key stage of animal growth and development, which determines the phenotypic characteristics of early and adult growth and development (L.Y. et al., 2016). Low nutrition intake of a maternal ewe can maximise the normal growth and development of the foetus; only when the ewe nutrition is seriously inadequate can it affect the growth and development of the foetus. Therefore, proper nutrition level plays an important role in the growth and development of lambs. In this study, for new-born and weaned lambs, all indexes in body size decreased at first and then increased with the increase of protein. In the body size at birth, there were differences or trends in body length, chest circumference and abdominal circumference. In the weaning body size, there were still differences or trends in body length, chest circumference and abdominal circumference. This difference may cause the growth and development of high-protein lambs to be better than low-protein and medium-protein lambs in the subsequent growth of lambs.

The evaluation of plasma indicators suggests accurate references to understand the results related to ruminant performance and nutritional status. In the current study, before delivery, different protein contents only affected the content of BUN, NH3L, CPRL3 and GLU. Low BUN content indicates high nitrogen metabolic efficiency (Bishonga et al., 1994). High blood urea levels indicate a high protein intake or excessive muscle mobilisation (Chimonyo et al., 2002). The BUN concentration of the high protein group was the highest among the three groups, and the body weight and weight gain of the ewes in the high protein group were better than those of the ewes in the low protein group and the middle protein group. At the same energy level, protein intake had a significant impact on the serum urea nitrogen content of lactating sheep, and with the decrease of protein intake, the serum urea nitrogen content showed a decreasing trend, which was consistent with Preston (1965). The ammonia produced by catabolism of various amino acids in various tissues of the body and the ammonia absorbed by the intestinal canal enter the blood to form blood ammonia. In this experiment, the NH3L content of ewes before delivery was higher than that before weaning, while the NH3L content of the high-protein group before delivery was the highest, indicating that the amino acid catabolism of high protein group was the fastest before delivery and had the greatest effect on the growth and development of its offspring lambs. CRPL3 is a kind of C-reactive protein, which is an indicator of infection. CRPL3 was the highest in the high protein group before delivery; therefore, low levels of protein should be used before delivery to avoid harm to ewes and lambs. Studies have shown that changes in serum biochemical indicators reflect the metabolism of such things as GLU serum content that can reflect the body’s energy metabolism (Graugnard et al., 2012). In this experiment, the level of GLU increased with an increase in dietary protein level, which was consistent with the results of Chelikani et al. Although dietary protein levels had no significant effect on ewe growth performance, lamb birth weight and litter weight, it did affect serum urea nitrogen and glucose, indicating that amino acid metabolism was affected along with glucose metabolism. Knowing that both the synthesis and excretion of urea are activities that involve energy expenditures, the energy metabolism and gluconeogenesis are relevant processes to take into consideration since they could be used in muscle deposition to promote the growth of the lamb (Noro and Wittwer,2012).

Before weaning the lamb, different protein levels only cause significant differences in TG. TG is the main lipid in animals and plants, and in serum, it reflects the body’s utilisation of lipids. The lower the TG, the higher the utilisation rate of fat and triglycerides in the blood are fuel sources (Caton et al., 2010). In this study, the highest TG content before weaning was 0.29 mmol/L (medium protein), and the low level and high protein group TG was 0.22 mmol/L. It indicates that the fat utilisation rate of low protein and high protein groups was higher. During the lactation period, there was no significant difference in blood biochemical indexes and growth performance among the three groups of lambs. According to the growth performance of lambs during lactation, it is concluded that the change of protein level in late pregnancy has little effect on the lactation period of ewes because it has no effect on the nutritional composition of milk and blood biochemical indexes of lambs. The urea nitrogen of ewe before weaning was not affected, and the level of blood triglyceride was affected, indicating that the protein level in late pregnancy may influence the fat metabolism of ewes during lactation. We know from Maas et al. (2009), that in ruminants during late pregnancy and lactation, increasing free fatty acids in plasma as a consequence of decreased insulin concentrations and fat mobilisation from the adipose tissue works as an alternative energy source. This may be one reason that ewes’ fat metabolism is affected. Moreover, why the blood glucose concentration during lactation is higher than that before giving birth can be explained by the retention of glucose from gluconeogenesis after fat mobilisation similar to the observations reported by Chilliard et al. (2000). This may also explain why fat metabolism is affected. The concentration of TG during lactation is lower than that during pregnancy, which may be due to the energy provided by TG catabolism for milk synthesis (Tanvi D. et al., 2016).

## 5 Conclusion

The protein levels of pregnant ewes ranged from 11.25% to 13.75%, which significantly affected the body height, body length, chest depth, chest circumference, straight crown hip length and curved crown hip length of the lamb at birth as well as the body length, chest circumference, head width and head length in the weaning body size of the lamb. The protein level of late pregnancy had no significant effect on the production performance of lactating ewes, nor did it affect the nutritional composition of milk and blood biochemical indexes of lambs, but there may be some effects on fat metabolism of ewes. And it has no effect on the weight of singleton, twins and triplets. Therefore, the optimum protein concentration for the growth of ewes and subsequent lambs in this experiment is 11.25%. In production, the proper dietary protein level for ewes during late gestation is beneficial in ensuring the development of Hu lambs.

## Acknowledgments

This work is supported by the College of Life Sciences of Hunan normal University and Hubei Zhiqinghe Animal Husbandry Company, and all experimental procedures related to the animals were performed according to protocols approved by the Animal Care Advisory Committee of Hunan Normal University, Changsha, Hunan, China. Wang Qiye and Wang Yancan helped to complete the experiment.

## AUTHOR CONTRIBUTIONS

The present experiment was originally designed by huansheng Yang and qiye Wang. The acquisition, analyses and interpretation of the data were done by all authors, and manuscript preparation was the primary responsibility of Xin Wang. All authors have read and approved of the submitted and revised versions of the paper.

## ANIMAL WELFARE STATEMENT

The author of the animal Welfare Statement confirms that the journal’s ethics policy, as noted on the journal’s author’s guide page, has been complied with and has been approved by the appropriate ethics review board. The authors confirm that they have followed EU animal protection standards and feed legislation for scientific purposes..

